# Hypusinated eIF5A is required for the translation of collagen

**DOI:** 10.1101/2021.02.23.432471

**Authors:** Marina Barba-Aliaga, Adriana Mena, Vanessa Espinoza, Nadezda Apostolova, Mercedes Costell, Paula Alepuz

## Abstract

The evolutionary conserved elongation factor eIF5A is required for the translation of mRNAs that encode protein sequences with consecutive prolines or combined with glycine and charged amino acids. Mammalian collagens are enriched in putative eIF5A-dependent Pro-Gly-containing tripeptides. Here, we show that eIF5A is needed for heterologous expression of collagen in yeast, and using a dual luciferase reporter system we confirmed that eIF5A depletion interrupts translation at Pro-Gly-collagenic motifs. Using mouse fibroblasts, we showed that depletion of active eIF5A reduced collagen 1α (Col1a1) content, which became concentrated around the nuclei, in contrast to a stronger and all over the cell collagen signal in untreated cells. Active eIF5A-depleted mouse fibroblast showed upregulation of endoplasmic reticulum (ER) stress markers, suggesting retention of partially synthesized Col1a1 in the ER. A dramatically lower level of Col1α1 protein was also observed in functional eIF5A-depleted human hepatic stellate cells treated with the profibrotic cytokine TGF-β1. Our results show that collagen expression requires eIF5A and imply its potential as a target for regulating collagen production in fibrotic diseases.

## Introduction

The translation elongation factor eIF5A, the eukaryotic homolog of prokaryotic elongation factor P (EF-P), is an essential and highly conserved protein. It is the only known protein modified by hypusination, which is required for eIF5A activity and occurs in two sequential enzymatic steps catalyzed by a deoxyhypusine synthase (DHPS) and a deoxyhypusine hydroxylase (DOHH), two enzymes that are also essential in most eukaryotic cells (Park & Wolff, 2018). In mammals, eIF5A is encoded by the paralogous genes *eIF5A1* and *eIF5A2*, whose amino acid sequences of the corresponding protein isoforms show high identity. Overexpression of each human isoform has been linked to different types of cancer, and high levels of eIF5A2 enhance metastasis (Mathews & Hershey, 2015; Nakanishi & Cleveland, 2016; Ning *et al*, 2020).

Hypusinated eIF5A binds to the ribosome in the E-tRNA site, where it interacts with the P-tRNA to promote productive positioning for peptide bond formation (Dever *et al*, 2018). Experiments with bacterial EF-P (Doerfel *et al*, 2013; Ude *et al*, 2013) and *Saccharomyces cerevisiae* (Gutierrez *et al*, 2013; Li *et al*, 2014) show that eIF5A facilitates translation through polyproline sequences containing three or more consecutive prolines, and this function seems conserved through evolution (Munoz-Soriano *et al*, 2017). Ribosome profiling (Schuller *et al*, 2017) and 5PSeq assays (Pelechano & Alepuz, 2017) have shown that eIF5A also alleviates ribosome pauses at tripeptide combinations of proline, glycine, and charged amino acids. A search in the human proteome identified that the “extracellular matrix (ECM)” compartment is highly enriched in potential eIF5A targets (Pelechano & Alepuz, 2017).

Collagen is the most abundant protein in vertebrates, constituting more than 25% of human body weight. Collagens are essential in the ECM and function in tissue structure, development and remodeling, cell adhesion and migration, cancer, and angiogenesis. The collagen family, which comprises around 28 different types in mammals, has characteristic triple-helical folding formed by three collagen α chains. In mammals, the more than 40 distinct α chains contain abundant repetitions on the so-called collagenic motif (X-Y-G), with proline or hydroxylated proline frequently in the first and second positions and glycine in the third, which enables the formation of the collagenous triple helix (Brodsky & Persikov, 2005; Kadler *et al*, 2007; Myllyharju & Kivirikko, 2001).

Collagen α chains are co-translationally inserted into the endoplasmic reticulum (ER) and are thus targeted to the secretory pathway. In the ER, the correct folding of the α chains to form procollagen requires the action of co- and post-translational modification enzymes, such as hydroxylases and glycosyltransferases, and the assistance of several molecular chaperones (Ito & Nagata, 2019). Secretion of the collagen triple helix involves the packaging and formation of big COPII vesicles (Malhotra & Erlmann, 2015). Incorrect folding or assembling of collagen results in the intracellular retention of immature procollagen and the induction of ER stress and the unfolded protein response (UPR) (Wong & Shoulders, 2019).

Deficient production of mature collagen can lead to severe diseases, collectively called collagenopathies (Arseni *et al*, 2018; Jobling *et al*, 2014; Myllyharju & Kivirikko, 2001), and to defects in wound healing when injured skin requires a physiological increase in collagen production (Nystrom & Bruckner-Tuderman, 2019). By contrast, excessive and uncontrolled synthesis of ECM proteins, especially collagen, results in fibrosis. Collagen I represents 80–90% of the proteins in the fibrotic matrix and disrupts normal tissue architecture and function. Fibrotic diseases, primarily liver fibrosis, followed by cardiac, renal, and pulmonary fibrosis, present a major health problem because of a lack of effective curative treatments and they are a leading cause of morbimortality (Ricard-Blum *et al*, 2018; Weiskirchen *et al*, 2019).

In the present study, we tested the hypothesis that the translation elongation factor eIF5A is necessary for collagen synthesis. First, we used yeast cells carrying temperature-sensitive eIF5A to determine levels of heterologously expressed Col1a1 and translation stops at collagenic tripeptide motifs cloned in a dual-luciferase system. Second, we used cultured mouse fibroblasts and depleted active eIF5A, either through treatment with the DHPS inhibitor GC7 (N^1^-guanyl-1,7-diaminoheptane) or with siRNA against DHPS and eIF5A1, to evaluate the expression of collagen type I α1 chain (Col1a1) and to localize it. We also explored the induction of ER stress upon depletion of active eIF5A. Third, we used TGF-β1 as a profibrotic stimulus in cultured human hepatic stellate cells (HSCs) to assess the extent to which Col1a1 overproduction depends on eIF5A expression. In yeast our data suggest that ribosomes stall when encountering collagenic motifs in eIF5A-depleted cells. Our results with mammalian cells show that active eIF5A is required to maintain Col1a1 homeostasis and to increase its production during fibrotic processes. We propose that eIF5A could be a therapeutic target for regulating collagen production in fibrotic diseases.

## Results

### Mammalian collagens, but not other ECM proteins, are enriched in putative eIF5A-dependent non-polyproline (PG)-containing tripeptides

Our previous study showed that the gene ontology terms “ECM organization” and “collagen metabolism” were significantly enriched in the human proteins with a high content (>25) of eIF5A-dependent motifs (Pelechano & Alepuz, 2017). To determine if only collagens or also other proteins of the mammalian ECM compartment contain a significant number of eIF5A-dependent tripeptides, and to identify which ones, we analyzed the amino acid sequence of the *Mus musculus* ECM proteins. ECM proteins are classified as fibrous proteins, proteoglycans, glycoproteins, and other proteins (Frantz *et al*, 2010), and we further subdivided fibrous proteins into collagens and others (Fig 1A). We examined the abundance of the 43 tripeptide motifs with the highest score for eIF5A-dependent ribosome pausing (Pelechano & Alepuz, 2017) and found that, although the average distribution of eIF5A-dependent motifs in mouse ECM proteins was 22.6 motifs per protein, collagens had a much higher average frequency (86.0 motifs per protein) than the rest of the ECM protein groups (motifs per protein: 21.7 in non-collagen fibrous proteins, 9.4 in proteoglycans, 13.0 in other proteoglycans, and 6.8 in others) (Fig 1B). Collagens also contained different eIF5A-dependent motifs from the rest of the ECM proteins, such as the non-polyproline motifs PGP, PPG, EPG, and DPG that constitute 78% of the motifs, whereas the well-known PPP (Dever *et al.*, 2018) motif was almost absent in collagen sequences (Fig 1B and 1C). Those four non-polyproline eIF5A motifs appear in stretches in the collagenic regions of the polypeptide sequences that form the triple helix of the collagen proteins (Kadler *et al.*, 2007). Our analysis points to mammalian collagens as important targets of eIF5A during their translation.

**Figure 1.**
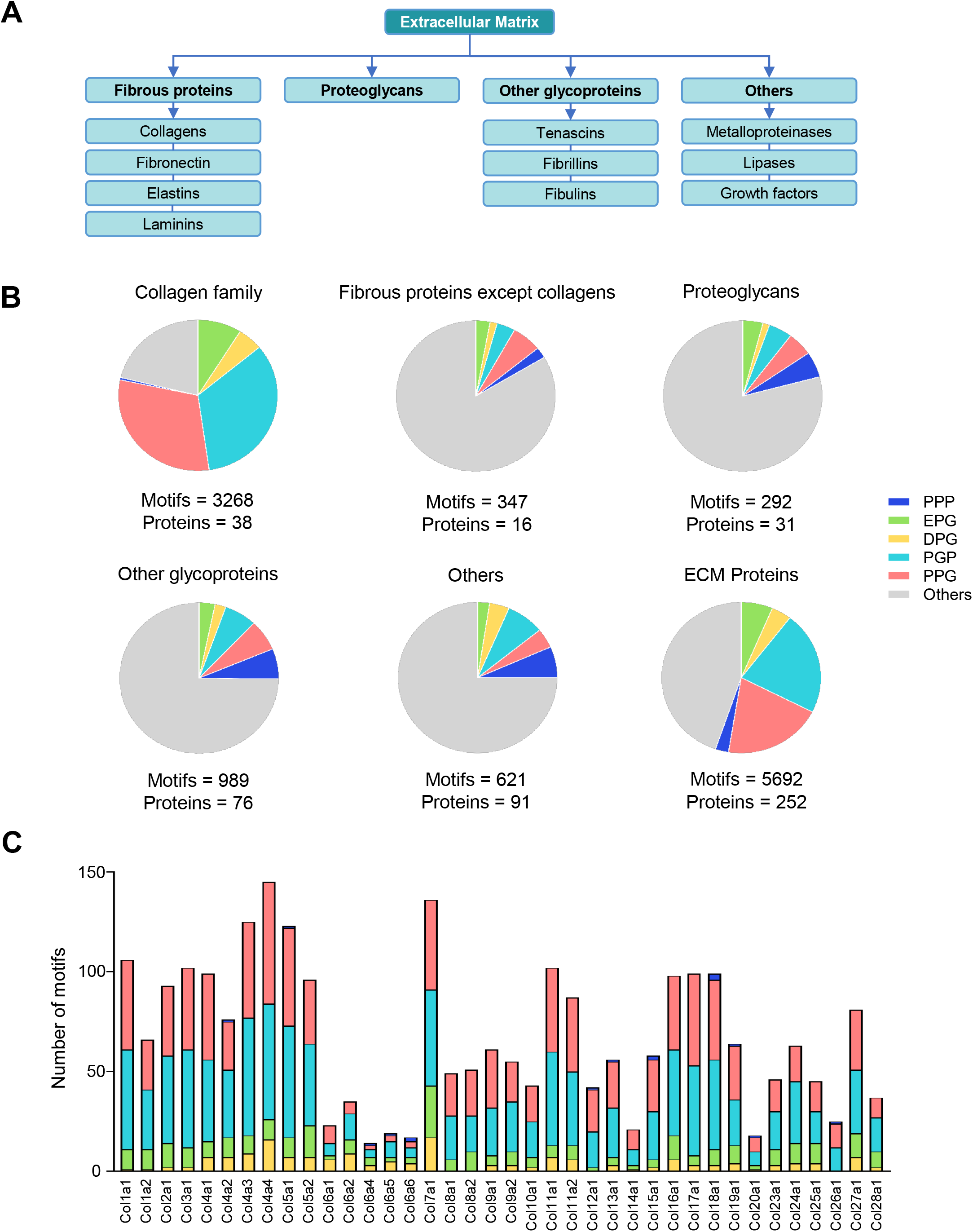
Distribution of eIF5A-dependent motifs in the mouse proteins of the extracellular matrix (ECM) A Classification of ECM proteins according to their structure and/or composition. B Distribution of the 43 highest-scoring eIF5A-dependent ribosome-pausing motifs (Pelechano & Alepuz, 2017) (polyproline PPP, collagenic EPG, DPG, PGP, PPG motifs, and others) in the subtypes of mouse ECM proteins. CNumber of eIF5A-dependent motifs in the mouse collagens.

### eIF5A stimulates translation of collagenic motifs expressed in yeast cells

To obtain molecular evidence supporting the role of eIF5A in the translation of collagenic sequences carrying putative elF5A-dependent motifs, we followed two different approaches using *Saccharomyces cerevisiae* cells. Human and yeast eIF5A proteins share >60% homology and are functionally interchangeable (Schnier *et al*, 1991; Schwelberger *et al*, 1993). Yeast cells do not contain collagen, but heterologous expression of human collagen has been successfully achieved (Brodsky & Ramshaw, 2017). First, using a tetracycline-regulated (tetO_7_) expression system (Gari *et al*, 1997) to avoid the deleterious effects of constant collagen expression in yeast cells, we expressed the first 420 amino acids of mouse Col1a1, containing 15 PPG, 10 PGP, and 3 EPG motifs (Fig 2A and Fig EV1A), fused to the β-galactosidase reporter. As a positive control of eIF5A-translation dependency, we constructed another fusion with a fragment of the yeast Bni1 protein containing three documented eIF5A-dependent polyproline sequences (Li *et al.*, 2014) (Fig 2A and Fig EV1A). We analyzed the expression of non-fused *lac*Z, Col1a1-*lac*Z, and Bni1-*lac*Z after inducing expression by incubating for 6 h in tetracycline-free media in wild-type yeast cells and eIF5A temperature-sensitive mutant cells (*tif51A-1*) carrying a single point mutation (Pro83 to Ser) (Li *et al*, 2011). β-galactosidase activity corresponding to Col1a1-*lac*Z and Bni1-*lac*Z was similar in wild-type and mutant cells when incubating cells at a permissive temperature (25°C) (Fig 2A). However, when incubated for 6 h at a restrictive temperature (37°C) (Li *et al.*, 2014), Col1a1-*lac*Z expression was reduced by ~40% in the mutant with respect to the wild type; a stronger reduction (~80%) was observed in Bni1-*lac*Z expression (Fig 2A). We did not observe any difference in the expression of non-fused *lac*Z between wild-type and mutant cells at 37°C (Fig 2A). These results indicate that eIF5A is required for the expression of the cloned Col1a1 fragment, and confirm that it is a strong requirement for the translation of the polyproline motifs of Bni1.

**Figure 2.**
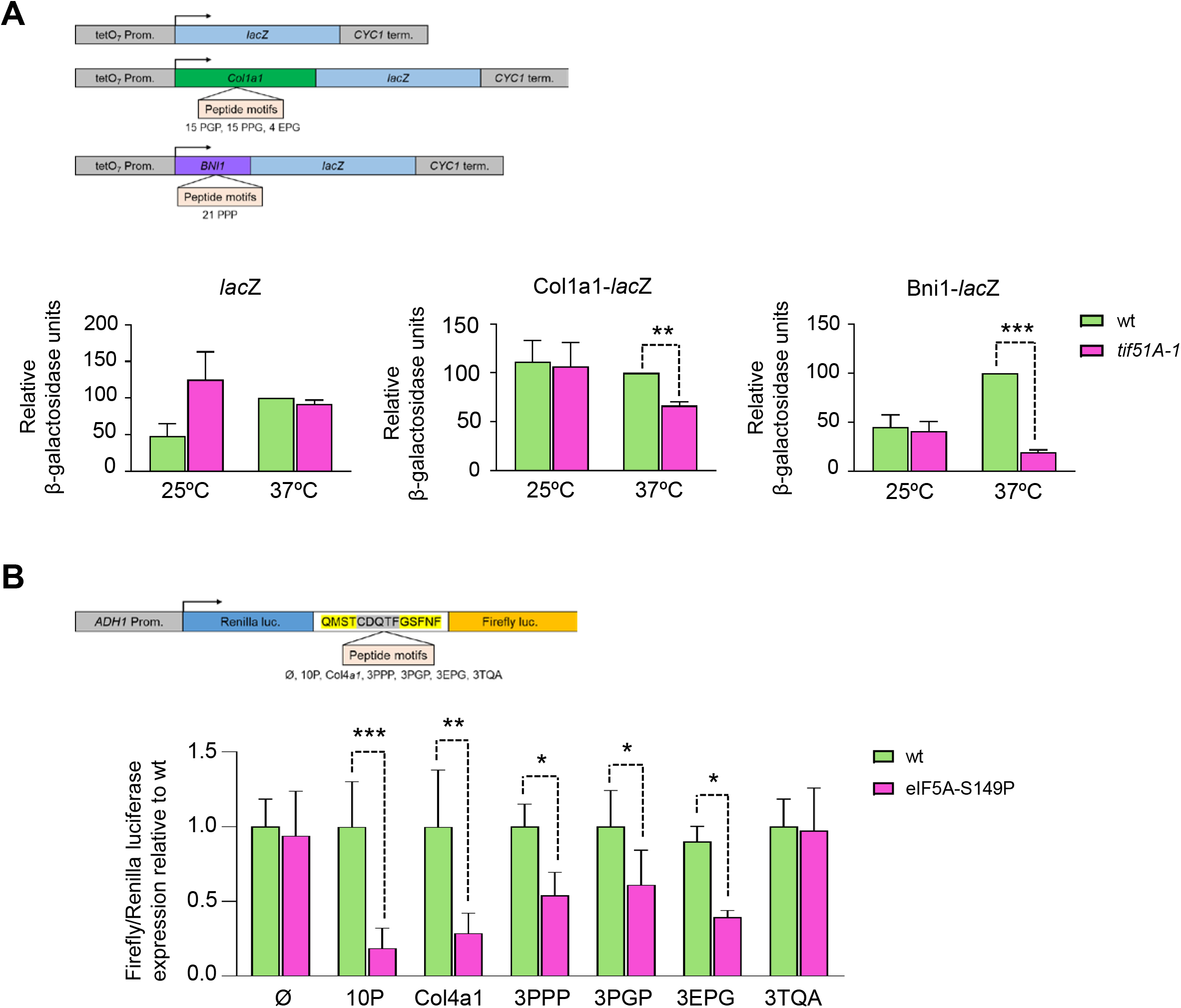
eIF5A stimulates translation of collagenic motifs in yeast cells. A Scheme of the lacZ expression plasmid (pCM179) and derivative plasmids containing fusions between a fragment of mouse collagen type I α1 chain (Col1a1) with collagenic motifs or a fragment of the yeast polyproline protein Bni1 and the *lac*Z gene expressed under the control of the doxycycline-regulated tetO_7_ promoter. These plasmids were introduced into isogenic stains expressing wild-type or temperature-sensitive eIF5A-P83S (*tif51A-1*), and β-galactosidase activity was assayed in a minimum of three independent experiments. Data are presented as the β-galactosidase units relative to the units of wild type at 37°C (given as 100 units). B Scheme of *Renilla*–firefly luciferase construct and peptide motifs inserted. Yellow and gray highlighted letters are flanking sequences to the insertion sites. Dual-luciferase reporter constructs were introduced into isogenic strains expressing wild-type eIF5A or temperature-sensitive eIF5A-S149P and grown at a semi-permissive temperature (33°C). The firefly:*Renilla* luciferase ratio for each construct is shown for a minimum of three independent experiments. A, B Data are presented as mean+standard deviation. Statistical significance was determined using a Student’s t-test relative to corresponding wild-type cells. *P<0.05, **P<0.01, ***P<0.001. See also Figure EV1.

To further define the role of eIF5A in the translation of specific collagenic motifs, we used a second approach based on the use of the dual-luciferase reporter system developed in yeast cells (Letzring *et al*, 2010). Dual-luciferase plasmids express a single mRNA that contains the *Renilla* luciferase at the 5′ end and the firefly luciferase open reading frames (ORFs) at the 3′ end, joined in-frame by a short sequence where the motifs to study can be cloned (Fig 2B and Fig EV1B). For these analyses we used a plasmid with no insertions; a plasmid with 10 consecutive optimal codons for proline inserted (10P) (Letzring *et al.*, 2010); and four plasmids we constructed with insertions of three non-consecutive repetitions of optimal codons for the polyproline motif PPP (3PPP), for the collagenic motifs PGP (3PGP) and EPG (3EPG), and for the control motif TQA (3TQA), for which ribosome pausing is not predicted upon eIF5A depletion (Pelechano & Alepuz, 2017) (Fig 2B and Fig EV1B). We also constructed a dual-luciferase plasmid in which we inserted a short stretch of mouse collagen IV sequence (Col4a1) containing several PPG and PGP motifs (Fig 2B and Fig EV1B). None of the resulting combinations of three amino acids in the inserted sequences, except the motif under investigation, were predicted to induce ribosome pausing by eIF5A depletion (Fig EV1C) (Pelechano & Alepuz, 2017). Dual-luciferase reporter constructs were introduced into isogenic strains expressing wild-type eIF5A or temperature-sensitive eIF5A-S149P and grown at a semi-permissive temperature (33°C) as described (Gutierrez *et al.*, 2013). We hypothesized that if eIF5A promoted translation of the inserted motifs in the dual-luciferase plasmid, then the firefly:*Renilla* luciferase ratio would be lower in the eIF5A-S149P mutant strain grown at 33°C. As shown in Figure 2B, the firefly:*Renilla* luciferase ratio for each construct was different depending on the cloned motif. The empty plasmid and the control TQA motif showed no difference in the firefly:*Renilla* luciferase ratio between the wild type and eIF5A mutant. As previously reported (Gutierrez *et al.*, 2013), a very low ratio was observed for the 10P construct upon eIF5A depletion (Fig 2B). Low ratios in the eIF5A mutant were also obtained with 3PPP, indicating that three non-consecutive polyproline tripeptide motifs are sufficient to impose a requirement for eIF5A, although without reaching the strong reduction observed with the longer polyproline motif (10P). Similarly, insertion of 3PGP and 3EPG and the Col4a1 fragment resulted in a lower luciferase ratio in the eIF5A mutant than in the wild type (Fig 2B). These results confirm that eIF5A promotes the translation of the collagenic tripeptide motifs.

### Depletion of functional eIF5A in mouse fibroblasts reduces collagen protein levels

To investigate the role of eIF5A in collagen synthesis in mammalian cells, we used a mouse fibroblast line. Fibroblasts are prototypical collagen producer cells, and we focused on Col1 because it is the most abundant collagen in animal tissue (Myllyharju & Kivirikko, 2001). Col1 is a heterotrimer with two α1 chains and one α2 chain. The Col1a1 subunit contains 50 PGP, 45 PPG, 10 EPG, and one DPG motif (Fig 1C). Using these mouse fibroblasts, we investigated the effect of inhibiting eIF5A hypusination with GC7, which acts as a competitive inhibitor of DHPS (Park & Wolff, 2018), and quantified Col1a1 levels. As seen in Fig 3A and 3B, treatment of fibroblasts with 65 or 30 μM GC7 reduced eIF5A hypusination and concomitantly lowered Col1a1 levels. A reduction in Col1a1 was already evident at 24 h of GC7 treatment and was still visible at 96 h. Col1a1 synthesis is mainly regulated at the level of mRNA stability and translation (Zhang & Stefanovic, 2016). To investigate whether the effects of eIF5A depletion were attributable to a negative effect on Col1a1 translation rather than transcription, we quantified mRNA levels after treatment with 30 μM GC7. No significant differences were observed in *Col1a1* mRNA levels with GC7 (Fig EV2A), suggesting that the effect of eIF5A inhibition on Col1a1 occurs at the level of translation. Because eIF5A is an essential protein in eukaryotic cells, we checked whether GC7 treatment reduced fibroblast viability. However, treatment with 30 μM GC7 up to 96 h did not significantly reduce fibroblast viability when compared to control (untreated) cells (Fig EV2B). Low cytotoxicity at 30 μM GC7 has also been described for human cells in culture (Xu *et al*, 2014); therefore, we used this GC7 concentration in the following experiments.

**Figure 3.**
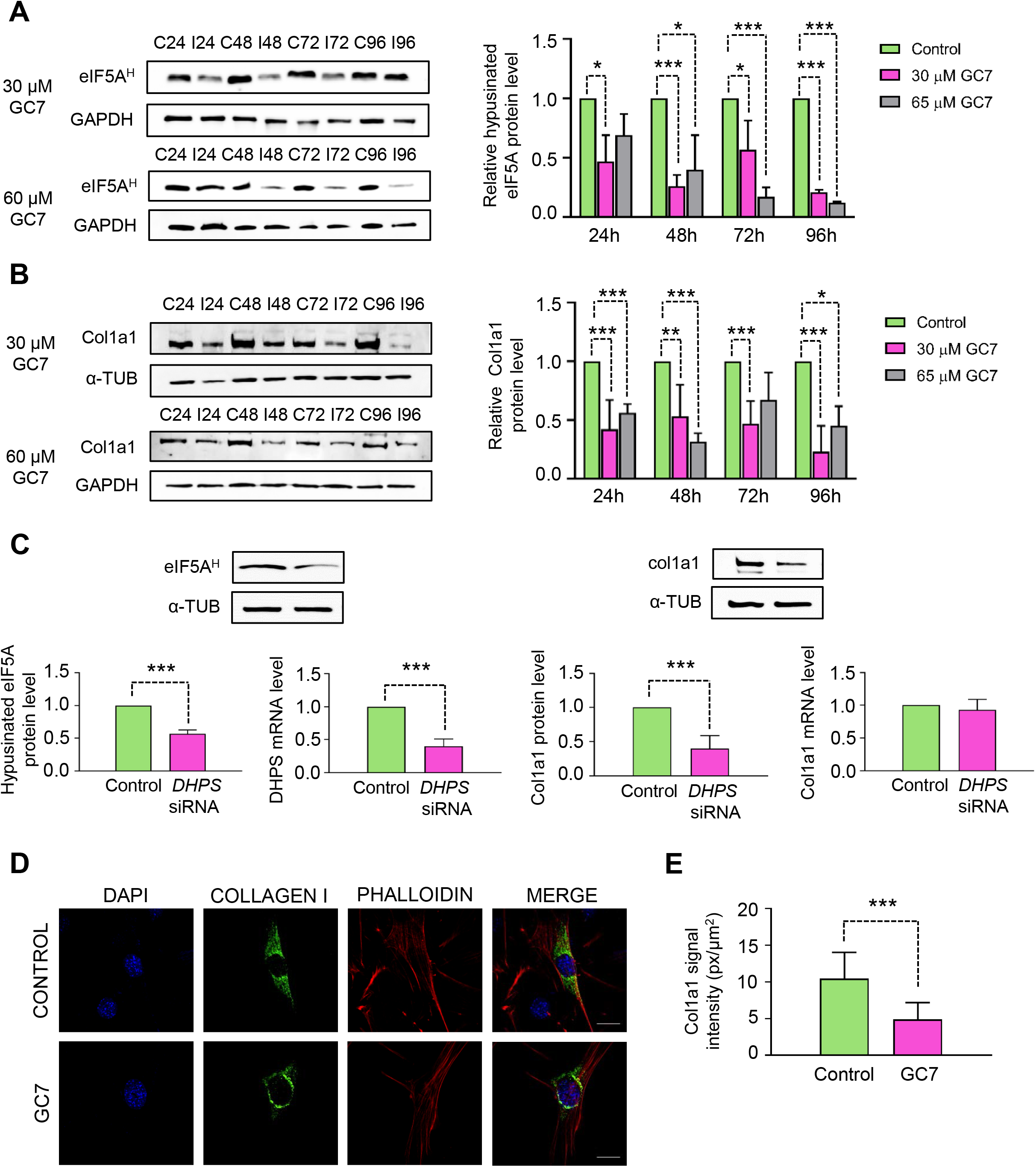
Depletion of functional eIF5A reduces collagen type I α1 chain (Col1a1) protein levels in mouse fibroblasts. A, B Western blotting and quantification analysis of hypusinated eIF5A (A) and Col1a1 (B) in fibroblasts at indicated time points after 30 μM or 65 μM GC7 treatment. GAPDH or α-tubulin protein levels were used as loading controls. A representative experiment is shown (n=6). Data are presented as mean+standard deviation (SD). Statistical significance was determined using a Student’s t-test relative to the corresponding untreated cells. *P<0.05, **P<0.01, ***P<0.001. C Western blotting, quantification, and RNA expression analysis of Col1a1, hypusinated eIF5A, and deoxyhypusine synthase (DHPS) after cell transfection with *DHPS* siRNA for 72 h. α-tubulin protein levels and *GAPDH* mRNA levels were used for normalization. A representative experiment is shown (n=5). Data are mean+SD. Statistical significance was determined using a Student’s t-test relative to scramble siRNA transfected cells. ***P<0.001. See also Figure EV2 and Figure EV3. D Confocal fluorescence microscopy images showing mouse fibroblasts at 96 h post-treatment with 30 μM GC7 stained with anti-collagen type I α1 chain (Col1a1) antibody (green), DAPI (blue), and phalloidin actin (red). Scale bars: 20 μm. See also Figure EV4 E Col1a1 intensity signal quantification from (D).

As an alternative to using GC7 to deplete functional eIF5A, we transfected mouse fibroblasts with *DHPS* siRNA to reduce the first enzymatic step of eIF5A hypusination. After 72 h, we observed a >50% reduction in *DHPS* mRNA levels and a similar reduction in hypusinated eIF5A (Fig 3C). This depletion of functional eIF5A correlated with a <50% reduction in Col1a1 protein levels, whereas no reduction in *Col1a1* mRNA was observed (Fig 3C). We then transfected mouse fibroblasts with *eIF5A1* siRNA to deplete *eIF5A1* mRNA, which is highly expressed in mammalian cells (Park & Wolff, 2018). The reduction in *eIF5A1* mRNA (~50% at 72 h of transfection) was paralleled by a strong reduction in Col1a1 protein (~70% compared to the control with scrambled siRNA) with, again, no change in *Col1a1* mRNA level (Fig EV3).

Together, these results show that hypusinated eIF5A is necessary to maintain high levels of Col1a1 protein in mouse fibroblasts and suggest a role for eIF5A in the translation of *Col1a1* mRNA.

### Depletion of functional eIF5A in fibroblasts leads to an accumulation of Col1a1 in perinuclear regions and ER stress

Translation of collagen polypeptides occurs at the ER where the polypeptide chains are translationally and post-translationally modified to form the triple helix (Malhotra & Erlmann, 2015; Sharma *et al*, 2017; Zhang & Stefanovic, 2016). To examine the effects of hypusinated eIF5A depletion on collagen synthesis, we analyzed Col1a1 distribution in mouse fibroblasts treated with 30 μM GC7. Immunostaining showed a lower Col1a1 signal in the treated fibroblasts than in untreated cells (Fig 3D and E and Fig EV4). Moreover, in the control cells the Col1a1 signal was visualized as dots concentrated more in the perinuclear region but also distributed all over the cytoplasm. By contrast, while the GC7-treated fibroblasts showed a concentration of Col1a1-stained dots in the perinuclear region, suggesting localization at the ER, the signal was almost absent in the rest of the cytoplasm (Fig 3D and Fig EV4).

It is well documented that misfolding and accumulation of mutated collagen at the ER lead to ER stress and the induction of collagenopathies (Gawron, 2016). Thus, if the lack of functional eIF5A is hindering the translation of the Col1a1 collagenic segments, it would stall translating ribosomes at the ER membrane and trigger the ER stress response. To test this, we analyzed the expression of several factors induced during the unfolded protein response that is activated in response to ER stress: the chaperone GRP78/BiP, a sensor of unfolded proteins in the ER; the ER stress transducer ATF6, a transcription activator of genes involved in protein folding, secretion, and degradation; and CHOP, a proapoptotic transcription factor induced upon severe ER stress (Hetz *et al*, 2020; Oyadomari & Mori, 2004; Ron & Walter, 2007). A quick and transient induction of *BiP*, *ATF6*, and *CHOP* mRNA levels was observed in fibroblasts after 24–48 h of GC7 treatment (Fig 4A). Correspondingly, CHOP protein levels increased around 100-fold at same time in GC7-treated fibroblasts (Fig 4B). Moreover, there was intense nuclear localization of CHOP protein in fibroblasts 24 h after GC7 treatment (Fig 4C and 4D); this correlates with its previously described change from cytoplasmic to nuclear localization during stress (Oyadomari & Mori, 2004; Ron & Walter, 2007).

**Figure 4.**
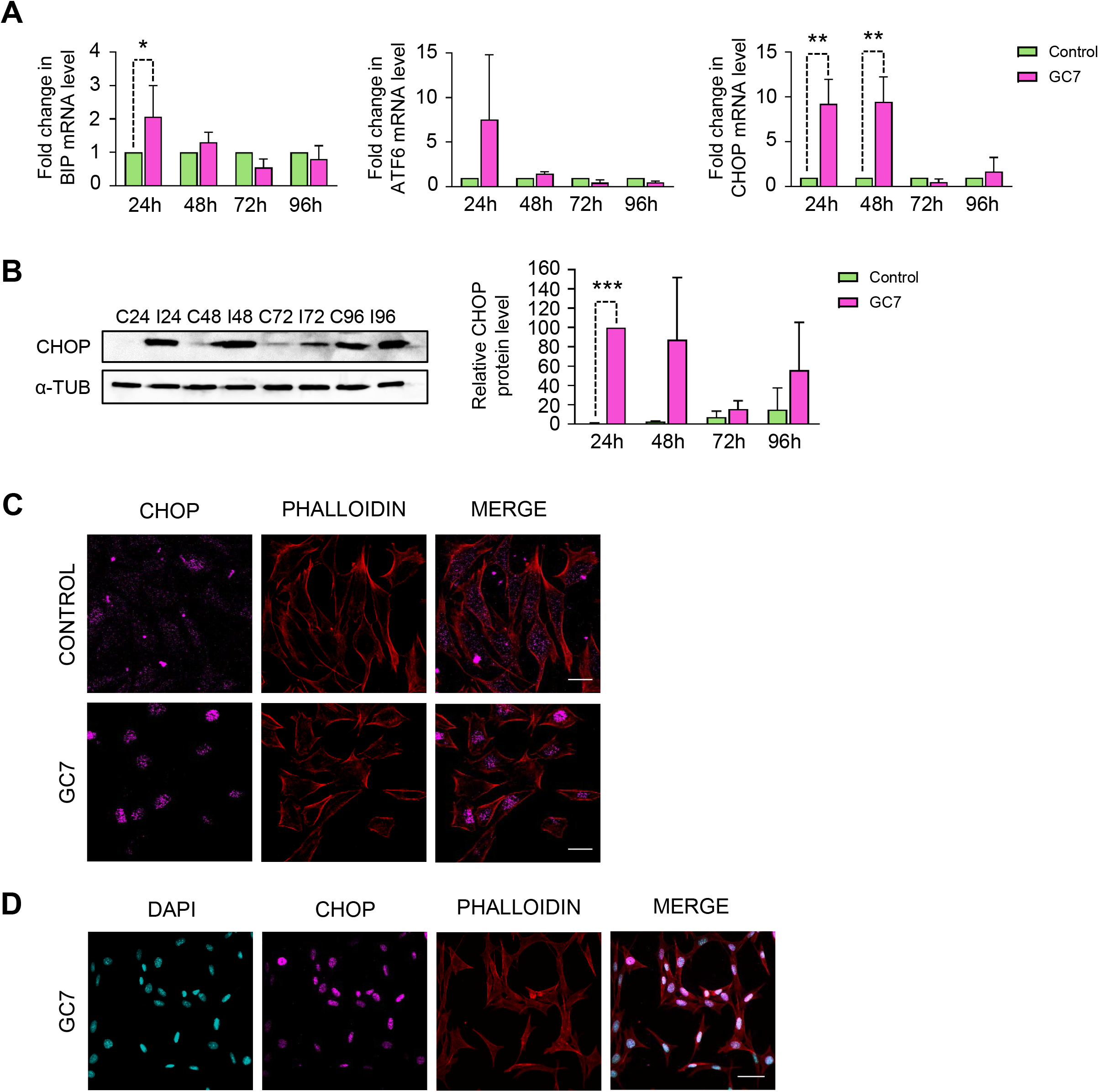
Depletion of functional eIF5A yields endoplasmic reticulum (ER) stress in mouse fibroblasts. A RT-qPCR analysis of the expression of the ER stress-related proteins BiP, ATF6, and CHOP at indicated time points after 30 μM GC7 treatment. *GAPDH* mRNA levels were used as an internal control. Data are presented as mean+standard deviation (SD) from five independent experiments. B Western blotting and quantification analysis of CHOP levels at indicated time points after 30 μM GC7 treatment. α-tubulin protein levels were used as loading control. A representative experiment is shown (n=3). Data are presented as mean+SD. C Confocal fluorescence microscopy images showing fibroblasts after 24 h of treatment with 30 μM GC7 stained with anti-CHOP antibody (magenta) and phalloidin actin (red). Scale bars: 20 μm. D Fluorescence microscopy images of fibroblasts treated as in (C) with DAPI (blue). Scale bars: 50 μm. A, B Statistical significance was determined using a Student’s t-test relative to corresponding untreated cells. *P<0.05, **P<0.01, ***P<0.001.

In sum, these results suggest that Col1a1 is retained at the ER in cells with a diminished amount of hypusinated eIF5A, causing a reduction in procollagen type I export from the ER to the Golgi apparatus and the induction of the unfolded protein response.

### Hypusinated eIF5A depletion inhibits in vitro TGF-β1-mediated fibrogenesis in human HSCs

Hepatic fibrosis is the most common fibrotic process, in which HSCs are the major cellular type responsible for producing high levels of collagen. HSCs in normal liver are quiescent and synthesize trace amounts of Col1, but an increase in the major profibrogenic inducer TGF-β1 (Meng *et al*, 2016) results in the transdifferentiation of HSCs into myofibroblast-like cells, which acquire a strong contractile phenotype and secrete and remodel large amounts of Col1 (De Minicis *et al*, 2007).

To assess the effect of eIF5A hypusination inhibition on Col1a1 production during hepatic fibrogenesis in vitro, LX2 cells, an immortalized cell line of human HSC, were treated with TGF-β1 (2.5 ng/mL) for 48 h, or with its vehicle (DMSO), together or not with 30 μM GC7. We observed an ~90% reduction in hypusinated eIF5A in GC7-treated cells compared to the corresponding controls of HSCs, both treated or not with TGF-β1 (Fig 5). The level of hypusinated eIF5A was higher in TGF-β1-treated than in control cells, but the difference was not statistically significant. Importantly, Col1a1 protein was undetectable in GC7-treated HSCs under basal conditions. As expected, a huge increase in Col1a1 levels was observed in TGF-β1-treated cells, which was fully prevented by co-treatment with GC7 (Fig 5). This result strongly suggests that eIF5A is required for Col1a1 synthesis during TGF-β1-mediated hepatic fibrogenesis.

**Figure 5.**
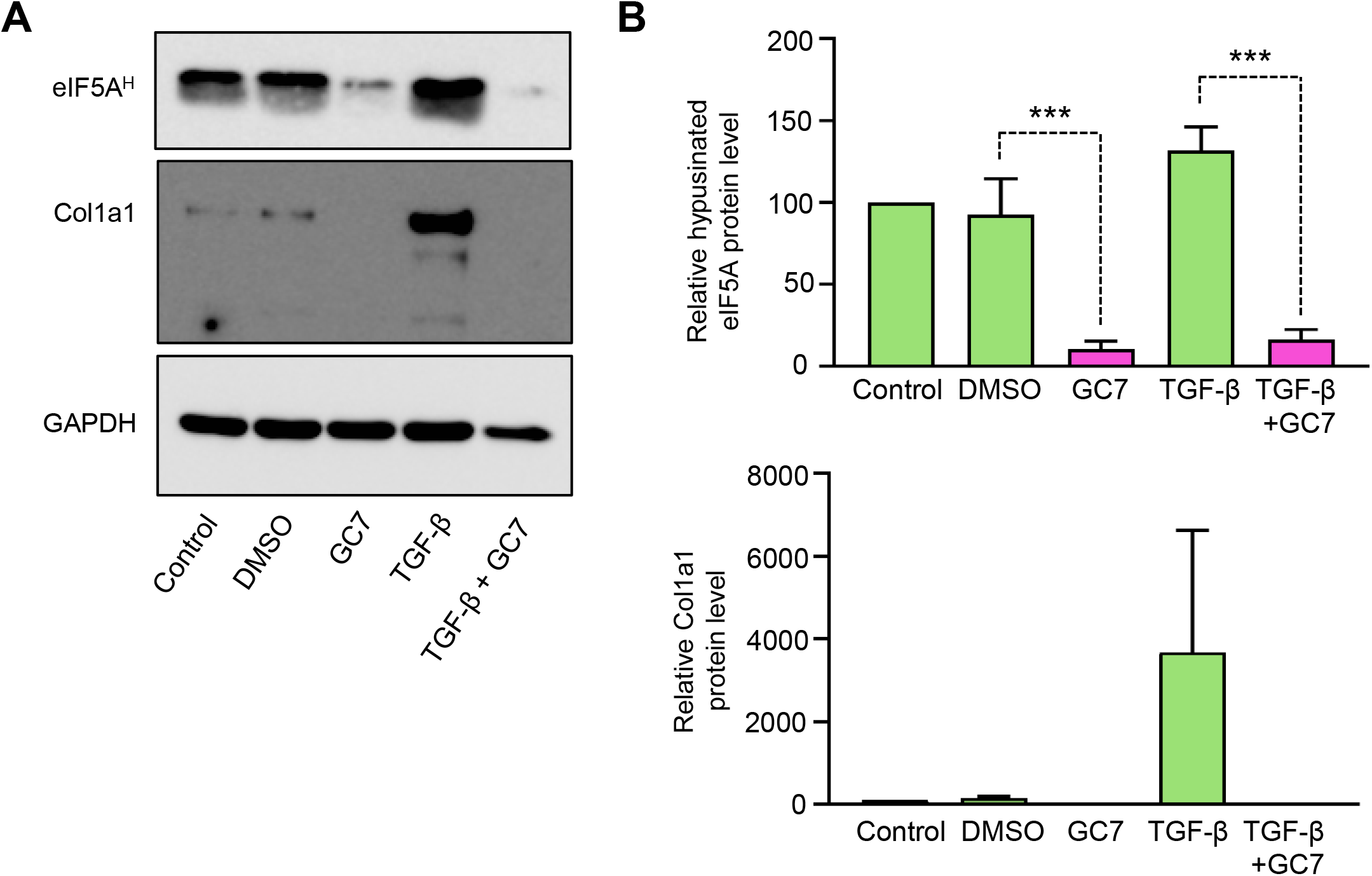
eIF5A is necessary for increased collagen production in cultured human hepatic stellate cells (HSCs) upon transforming growth factor-β1 (TGF-β1)-mediated fibrogenesis. A Western blotting analysis of collagen type I α1 chain (Col1a1) and hypusinated eIF5A in LX-2 human HSCs untreated (control) or treated with 30 μM GC7 (GC7) or vehicle (DMSO) for 48 h. TGF-β1 (2.5 mg/mL) was used as profibrogenic stimulus. GAPDH protein levels were used as loading control. A representative experiment is shown (n=3). B Quantification analysis of experiments in (A). Data are presented as mean+standard deviation. Statistical significance was determined using a Student’s t-test relative to corresponding vehicle or TGF-β1-treated cells. ***P<0.001.

## Discussion

Expression of collagen, the most abundant protein in animals, is regulated at the level of transcription, mRNA stability, and translation (Lindquist *et al*, 2000). Our results here, using different approaches, provide evidence that eIF5A is essential for the translation of tripeptide motifs that are heavily present in collagens. In yeast, fusions of mouse Col1a1 fragments with β-galactosidase show low expression levels upon eIF5A depletion. In a dual-luciferase reporter system, PGP, PPG, and EPG non-polyproline tripeptide motifs, abundant in collagens, stopped translation between *Renilla* and firefly luciferases when eIF5A was lacking. In mouse fibroblasts, depleting functional eIF5A by treatment with GC7 or by interfering with the *eIF5A1* or *DHPS* mRNAs results in reduced Col1a1 levels, with no reduction in the *Col1a1* mRNA. In human HSCs, inhibiting eIF5A hypusination with GC7 eliminates the Col1a1 overproduction caused by profibrotic treatment with TGF-β1. Together, these results link eIF5A activity with translation of the PGP, PPG, and EPG motifs that are highly abundant in the Col1a1 collagenic regions. The need for eIF5A in translating these non-polyproline tripeptide motifs has been suggested in yeast by ribosome profiling (Schuller *et al.*, 2017) and 5PSeq assays (Pelechano & Alepuz, 2017), in which it was noted that eIF5A depletion resulted in ribosomes stalling in these motifs. However, no physiological confirmation of a stop to translation causing the deficient protein synthesis in these motifs has been shown until now. Although our work has focused on the study of Col1a1, the existence of similar collagenic stretches in all collagens (Fig 1) suggests a similar dependency on eIF5A for the translation of the different collagens. Supporting this, a short polypeptide stretch of mouse Col4a1 containing several PPG and PGP motifs clearly stops translation in a yeast dual-luciferase system (Fig 2 and Fig EV1).

We observed that inhibition of eIF5A hypusination led to an accumulation of Col1a1 around the nuclei of mouse fibroblasts and provoked ER stress. Mutations in collagen or collagen-maturation enzymes or proteins that inhibit proper collagen folding, packing, and secretion to induce ER stress and intracellular retention of collagen have been observed in collagenopathies such as skeletal chondrodysplasia, osteogenesis imperfecta, and osteoarthritis (Gawron, 2016). Intracellular retention of procollagen induces the ER stress-proteins BiP and CHOP, which may lead to apoptosis (Schulz *et al*, 2016). eIF5A has also been implicated in ER function. In yeast and HeLa cells, depletion of eIF5A causes ER stress and upregulates stress-induced chaperones (Mandal *et al*, 2016). We propose a direct role of eIF5A in facilitating ER-coupled collagen translation; similarly, eIF5A may facilitate the co-translational translocation into the ER of other proteins containing eIF5A-dependent motifs. Although most collagenopathies have a genetic etiology, with a mutation in a collagen or collagen-metabolism protein (Forlino & Marini, 2016; Jobling *et al.*, 2014; Marini *et al*, 2017), it would be of interest to determine if patients with a clinical but not a genetic diagnosis have defects in eIF5A expression.

The excessive and uncontrolled synthesis of ECM proteins, mainly collagen type I, is the hallmark of fibrotic diseases; thus, the severity of the fibrotic disease depends on the amount of Col1 produced (Zhang & Stefanovic, 2016). Nearly 45% of all deaths in the developed world are attributed to chronic fibroproliferative diseases. Therefore, the demand for effective antifibrotic drugs will likely continue to increase in the coming years (Wynn, 2008). In our study, using an in vitro model of fibrogenesis with human HSCs, TGF-β1 induced the expected huge increase in Col1a1 levels, which was almost completely abolished upon GC7 treatment. In vivo studies are required to confirm the profibrotic role of eIF5A and test whether downregulation of eIF5A may be beneficial for the treatment of fibrosis of the liver and other organs.

## Materials and Methods

### Yeast strains, culture conditions, and plasmids

*S. cerevisiae* strains used are listed in Appendix Table S1. Yeast cells were grown in YPD (2% glucose, 2% peptone, 1% yeast extract) or synthetic complete media (SC) lacking the indicated amino acid (2% glucose, 0.7% yeast nitrogen base (YNB) and required Drop-Out percentage). For experiments, cells were grown exponentially until they reached the required OD_600_ at 25°C, when they were transferred to a semi-permissive (33°C) or restrictive (37°C) temperature.

Fusions with *Lac*Z and under the control of a tetracycline-repressible operon (tetO_7_) were constructed in the plasmid pCM179 (Gari *et al.*, 1997). Dual-luciferase reporter constructs were generated by cloning between the in-frame *Renilla* and firefly luciferase ORFs in the plasmid pDL202 (Letzring *et al.*, 2010). Inserts were generated by PCR amplification using *M. musculus* cDNA or *S. cerevisiae* genomic DNA. pDL202 and 10P plasmids were kindly supplied by Elizabeth Grayhack (University of Rochester, USA) (Letzring *et al.*, 2010). Oligonucleotides used are listed in Appendix Table S2. The GAP-REPAIR homologous recombination method was used for cloning (Joska *et al*, 2014).

### β-galactosidase assay in yeast

Wild-type eIF5A (BY4741) or temperature-sensitive eIF5A-P83S (*tif51A-1*) containing pCM179 and derivative plasmids were cultured in SC-URA medium at 25°C with 2 μg/mL doxycycline to keep the tetO_7_ promoter switched off. At an OD_600_ of 0.2, the cell culture was washed with medium lacking doxycycline, resuspended in fresh SC-URA medium to activate tetO_7_ transcription, and incubated at 25°C or 37°C for 6 h. Then, β-galactosidase assays were performed as described previously (Garre *et al*, 2012).

### Dual-luciferase assay in yeast

The dual-luciferase assay with yeast strains containing wild-type eIF5A (J697) or temperature-sensitive eIF5A-S149P (J699) were performed as described previously (Gutierrez *et al.*, 2013) with some modifications. Yeast was cultured in SC-URA medium at 33°C to an OD_600_ of 0.8, harvested, and frozen. Subsequently, cell pellets were resuspended in 200 μL of lysis buffer (50 mM Tris-HCl, pH 7.5, 150 mM NaCl, 5 mM MgCl_2_, 5% NP-40, cOmplete Protease Inhibitor Cocktail (Roche)) and mixed with one volume of glass beads for further homogenization in a Precellys tissue homogenizer (Bertin Corp.). Lysates were cleared by centrifugation at 12000 rpm for 5 min at 4°C and protein extracts were assayed for firefly and *Renilla* luciferase activity sequentially in a 96-well luminometry plate (Thermo Fisher Scientific) using a microplate luminometer (Thermo Fisher Scientific) and the Dual-Glo Luciferase Assay System (Promega). Finally, the firefly:*Renilla* luciferase ratio for each construct was calculated.

### Fibroblast cell line and culture

The fibroblast cell line was isolated from mouse kidney and immortalized by retroviral delivery of the SV40 large T (Benito-Jardon *et al*, 2017). These cells were cultured in DMEM containing 10% FCS (Thermo Fisher Scientific) and 1% penicillin-streptomycin (Gibco), and maintained at standard conditions of 37°C and 5% CO_2_ in a humidified atmosphere. Cells were cultured with or without treatment with the DHPS inhibitor GC7 (Calbiochem) at the indicated concentration to deplete functional eIF5A (Park *et al*, 1994).

### Hepatic stellate cell line and culture

LX2 (immortalized cell line of HSCs) were gifted by Dr. Scott L. Friedman, Icahn School of Medicine at Mountain Sinai, USA. Cells were cultured in DMEM with high glucose (Sigma-Aldrich), supplemented with 10% FCS, 2 mM L-glutamine, 50 U/mL penicillin, and 50 μg/mL streptomycin (Gibco, Invitrogen) in a humidified atmosphere with 5% CO_2_ at 37°C. For the experiment, cells were seeded in a 6-well plate (0.18 × 10^6^ cells/well) 24 h before treatment and treated for 48 h with TGF-β1 (2.5 ng/mL) or its vehicle DMSO (Sigma-Aldrich). GC7 was employed at 30 μM alone or together with TGF-β1.

### Fibroblast viability measurements

Non-treated cells and 30 μM GC7-treated cells were grown in DMEM media and samples taken at 10, 24, 48, 72, and 96 h. Cells from suspension, PBS washes, and the fibroblast monolayer detached with trypsin were collected for analysis. A 4% trypan blue/PBS dilution was used to stain dead cells. The fibroblast population was analyzed by counting live and dead cells in a Neubauer chamber. The cell death percentage was calculated from the 50–400 cells counted.

### Silencing DHPS and eIF5A expression by siRNA in fibroblasts

A synthetic pool of siRNAs was used to target and knock down DHPS (Sigma-Aldrich, #EMU150671) or eIF5A (Santa Cruz, #sc-40560) expression according to the manufacturer’s protocol. Briefly, 3×10^4^ cells/well were seeded in a 6-well culture plate 1 day prior to transfection until 70–90% confluent. For DHPS silencing, 250 μL Opti-Mem I Reduced Serum Medium (Thermo Fisher Scientific, #31985062) containing 7 μL of lipofectamine 3000 reagent (Thermo Fisher Scientific, #L3000015) and 10 μL of either *DHPS* or control siRNA (Sigma-Aldrich, #SIC001), previously incubated together for 15 min, was added to the culture. Cells were transfected and incubated for 72 h. For eIF5A silencing, the culture medium was replaced with transfection medium (Santa Cruz, #sc-36868) containing 6 μL of transfection reagent (Santa Cruz, #sc-29528) and 12 μL of either *eIF5A* or control siRNA (Santa Cruz, #sc-36869), previously incubated together for 30 min. The cells were transfected and incubated with the transfection mixture for 6 h, after which the transfection medium was replaced with 2× fibroblast usual medium. After 24 h, the medium was replaced with 1× usual medium and incubated until 72 h post-transfection. Finally, cells were collected for RT-qPCR and western blot analysis.

### Western blot analysis

For western blot analysis of fibroblasts protein content, proteins were extracted in RIPA buffer (150 mM NaCl, 0.05 M Tris-HCl pH 8.0, 0.005 mM EDTA, 0.1% SDS, 0.5% sodium deoxycholate, 1% NP-40, cOmplete Protease Inhibitor Cocktail (Roche)) for 15 min. The lysates were centrifuged at 13000 rpm at 4°C for 5 min to remove the cell debris and insoluble proteins. Protein content was quantified in a Nanodrop device (Thermo Fisher Scientific) and 100 μg of each sample was loaded into the 4–20% gradient precast SDS-PAGE gels (BioRad).

To analyze protein expression in LX2 cells, whole-cell protein extracts were obtained by lysis of the cell pellets in complete lysis buffer (20 mM HEPES pH 7.4, 400 mM NaCl, 20% (v/v) glycerol, 0.1 mM EDTA, 10 μM Na_2_MoO_4_, 10 mM NaF). Immediately prior to their use, 1 mM DTT, 5 mM broad-spectrum serine, cysteine protease inhibitors (Complete Mini™ and Pefabloc®, Roche Life Science), and 0.05% detergent solution (NP-40 Surfact-Amps™, Thermo Fisher Scientific) were added. Samples were then vortexed twice at maximum speed (10 s), incubated on ice (15 min), vortexed again at maximum speed (30 s), and centrifuged (16000 g, 15 min, 4°C).

Western blot membranes were incubated with primary antibodies against Col1 (rabbit polyclonal 1:500, Millipore Cat# AB765P, for fibroblasts and 1:1000 Cell Signaling Technology Cat# 84336, for LX2 cells), hypusinated eIF5A (FabHpu antibody, 1:600, Genentech), Gadd153 (anti-CHOP antibody, 1:200, Santa Cruz Biotechnology Cat# sc-7351), α-tubulin (1:5000, Proteintech Cat# 66031-1-Ig, RRID:AB_11042766), or GAPDH (mouse monoclonal anti-GAPDH antibody, 1:5000, Thermo Fisher Scientific Cat# AM4300, for fibroblasts and rabbit monoclonal Sigma-Aldrich Cat# G9545, 1:10000 for LX2 cells). Appropriate HRP-conjugated secondary antibodies were used (1:10000 Promega). Chemiluminescent signals were detected and digitally analyzed, and normalized against α-tubulin or GAPDH.

### mRNA analysis by RT-qPCR

To analyze mRNA levels, total RNAs were isolated from fibroblasts using the PureLink RNA Mini Kit (Invitrogen). The reverse transcription and quantitative PCR reactions were performed as detailed previously (Garre *et al*, 2013). Specific primers are listed in Supplementary Table 2. *GAPDH* mRNA levels were used for normalization.

### Immunofluorescence assays in fibroblasts

For indirect immunofluorescence of fibroblasts, cells (10^3^) were seeded onto laminin-coated glass coverslips and cultured for 1 or 4 days at 37°C in medium containing 30 μM GC7. Cells were fixed with 4% PFA for 10 min, washed four times with PBS, permeabilized with 0.1% Triton X-100 in PBS for 10 min, and blocked for nonspecific protein binding with 3% BSA/PBS for 1 h at room temperature. Cells were incubated with primary antibodies against Col1 (1:250, Merck #AB765P) or Gadd 153/CHOP (1:40, Santa Cruz #sc-7351) (in 1% BSA/PBS), and secondary antibodies Alexa Fluor 488-conjugated anti-rabbit (1:2000) or Alexa Fluor 546-conjugated anti-mouse (1:800) IgG. Rhodamine-conjugated phalloidin coupled with TRITC (1:500, Sigma-Aldrich) was used for actin filament counterstaining (in 1% BSA/PBS). Nuclei were counterstained with DAPI (1:10000, Merck). Immunofluorescence images were acquired using an Olympus FLUOVIEW FV 1000 confocal microscope equipped with ×40 and ×60 objectives on a Nikon ECLIPSE E200. The same exposure times were used to acquire all images. Image analysis was carried out with ImageJ and Photoshop CC 2019. To quantify collagen fluorescence intensity, the Col1a1 intensity signal was expressed relative to the area of each cell. ImageJ (SCR_003070) software was used to quantify control (n=21) and 30 μM GC7-treated cells (n=38) corresponding to 20 fields from two independent experiments.

### Statistical analysis

Statistical evaluation was carried out with Microsoft Office Excel 2016. Data are expressed as the mean±SD. Significant differences between the treated groups and the control were determined using Student’s t-test (two-tailed), at a significance level of P<0.05.

## Data availability

This study includes no primary dataset deposited in external repositories.

## Acknowledgements

P.A. was funded by the Ministerio Español de Economía y Competitividad (BFU2016– 77728-C3–3-P) and by the Regional Valencian Government (AICO2020/086). N.A. was funded by the Ministerio de Ciencia, Innovación y Universidades (RTI2018-096748-B-I00). M. B.-A. is a recipient of a predoctoral research contract (FPU2017/03542) from the Spanish Ministry of Science, Innovation and Universities. We thank Thomas Dever for his useful comments and suggestions. We are grateful to Irene Gimeno-Lluch for her initial input and to Sheila Otega-Sanchís for technical help with the fibroblast experiments. We are also thankful to all GFL laboratory members for their comments and support.

## Author contributions

P.A. conceived the study. M.B.-A., A.M., V.E., N.A., M.C., and P.A. acquired and analyzed the data. A.M., M.B.-A., V.E., N.A., M.C., and P.A. contributed to the methodology. P.A. and M.B.-A. drafted the manuscript. M.B.-A., A.M., V.E., N.A., M.C., and P.A. edited the manuscript.

## Expanded View Figure legends

**Figure EV1. Description of the constructs to study the role of eIF5A in the translation of collagenic motifs in yeast cells**

A Mouse Col1a1 and yeast protein Bni1 protein sequences cloned into pCM179 plasmids to measure expression of the lacZ fusions. In both sequences, collagenic motifs are in red.

B Sequences containing the tripeptide collagenic motifs cloned into the double luciferase reporter system (pDL202). Yellow and gray highlighted letters are flanking sequences to the insertion sites; red letters are polyproline (10P, 3PPP), collagenic motifs (3PGP, 3EPG), a mouse collagen IV fragment (Col4α1), and control motif (3TQA); blue letters are spacer sequences between motifs.

C Lateral tripeptide combinations due to the insertions do not produce pausing on the 5PSeq assay. Ribosome-pausing values obtained from 5PSeq assay (Pelechano & Alepuz, 2017) for the different tripeptide combinations of the sequences that result from cloning into the double luciferase reporter system. The ribosome pause value was obtained in a previous 5PSeq assay (Pelechano & Alepuz, 2017) and is indicated as log_2_ of the obtained signal for the temperature-sensitive eIF5A strain (eIF5Ats) relative to the wild-type eIF5A strain at restrictive temperature (37°C). The value is shown at the −17, −14, −11 and −8 positions upstream of the first nucleotide of the codon at the A position in the ribosome. Cut-off values were established as log2foldchange eIF5Ats/wt>2 and are shown in red.

**Figure EV2. Depletion of functional eIF5A in fibroblasts does not immediately decrease cell viability or Col1a1 mRNA levels**

A qPCR analysis of expression of *Col1a1* at indicated time points after 30 μM GC7 treatment. *GAPDH* mRNA levels were used as an internal control. Data are mean+SD from five independent experiments. Statistical significance was determined using a Student’s t-test relative to corresponding untreated cells. No statistically significant difference was found.

B Mouse fibroblast dead cell percentage at indicated time points after 30 μM GC7 treatment. Data are mean+standard deviation (SD) from six independent experiments. Statistical significance was determined using a Student’s t-test relative to corresponding untreated cells. **P<0.01.

**Figure EV3. siRNA depletion of eIF5A reduces Col1a1 protein levels in fibroblasts**

A, B Mouse fibroblast cells were transfected with *eIF5A-1* siRNA for 72 h. *eIF5A-1* (A) and *Col1a1* (B) mRNA levels were quantified from three independent experiments.

C Col1a1 protein expression was analyzed by Western blot using α-tubulin protein detection for normalization (n=4) and a representative experiment is shown (D).

Data are presented as mean+standard deviation. Statistical significance was determined using a Student’s t-test relative to scramble siRNA transfected cells. **P<0.01; ***P<0.001.

**Figure EV4. Depletion of functional eIF5A reduces collagen expression in mouse fibroblasts**

A Confocal fluorescence microscopy images showing mouse fibroblasts at 96 h post-treatment with 30 μM GC7 stained with anti-collagen type I α1 chain (Col1a1) antibody (green), DAPI (blue), and phalloidin actin (red). Scale bars: 40 μm.

## Appendix

**Appendix Table S1. Yeast strains used in this study**

**Appendix Table S2. Oligonucleotides used in this study**

