## Supplementary Figures 1-4 for "Hypusinated eIF5A is required for the translation of collagen"

**A**

**mCol1α1 (1-420 aa)**

MFSFVDLRLLLLLGATALLTHGQEDIPEVSCIHNGLRVPNGETWKPEVCLICICHNGTAVCDDVQCNEELDCPNPQRREGECCAFCPE
EYVSPNSEDVGVEGPKGD**PGP**QGPRGPVG**PPG**RDGIPGQPGL**PGPPGPPGPPGPPG**LGGNFASQMSYGYDEKSAGVSV**PGP**MGP
SGPRGL**PGPPG**A**PGP**QGFQG**PPGEPGEPG**GSGPMGPRG**PPGPPG**KNGDDGEAGKPGRPGERG**PPGP**QGARGLPGTAGLPGMKG
HRGFSGLDGAKGDAGPAGPKG**EPG**SPGENGAPGQMGPRGLPGERGR**PGPPG**TAGARGNDGAVGAAG**PPGP**TGPTG**PPG**FPGAV
GAKGEAGPQGARGSEGPQGVRG**EPGPPGP**AGAAGPAGNPGADGQPGAKGANGAPGIAGAPGFPGARGPSGPQGPSGPPGPKG

**yBni1 (1221-1357)**

NTDGAEDLSTQSSVLSSQ**PPPPPPPPPP**VPAKLFGESLEKEKKSEDDTVKQETTGDSPA**PPPPPPPPPPPP**MALFGKPKGET**PPPPP**
LPSVLSSSTDGVIPPAPPMMPASQIKSAVTSPLLPQSPSLFEKYPRPHKK

**B**

**pDL202:** QMSTCDQTFGSFNF

**pDL202-10Pro:** QMST**PPPPPPPPPP**GSFNF

**pDL202-Col IV:** QMSTAAAAAC**PPGFTGPPGPPGPPGPPG**YRCDQTFGSFNF

**pDL202-3PPP:** QMSTAAAAAC**PPP**YAC**PPP**YAC**PPP**YRCDQTFGSFNF

**pDL202-3PGP:** QMSTAAAAAC**PGP**YAC**PGP**YAC**PGP**YRCDQTFGSFNF

**pDL202-3EPG:** QMSTAAAAAC**EPG**YAC**EPG**YAC**EPG**YRCDQTFGSFNF

**pDL202-3TQA:** QMSTAAAAAC**TQA**YAC**TQA**YAC**TQA**YRCDQTFGSFNF

**C**

Cloned sequence: AAC**PPP**YAC**PPP**YAC**PPP**YR

| Ribosome pausing 5Pseq (log2 Fold change eIF5Ats/wt) | | | | |
|---|---|---|---|---|
| **Motif** | **Ribosome position** | | | |
| | **-17** | **-14** | **-11** | **-8** |
| AAC | 0.81 | 0.85 | 0.90 | 0.94 |
| ACP | 1.07 | 0.97 | 0.85 | 0.98 |
| CPP | 1.16 | 0.81 | 1.32 | 1.13 |
| **PPP** | 1.20 | 1.78 | **2.56** | 1.59 |
| PPY | 1.33 | 1.54 | 1.57 | 1.31 |
| PYA | 1.47 | 1.27 | 1.53 | 1.10 |
| YAC | 0.94 | 1.13 | 0.92 | 1.18 |
| PYR | 1.03 | 0.78 | 1.11 | 0.97 |

Cloned sequence: AAC**PGP**YAC**PGP**YAC**PGP**YR

| Ribosome pausing 5Pseq (log2 Fold change eIF5Ats/wt) | | | | |
|---|---|---|---|---|
| **Motif** | **Ribosome position** | | | |
| | **-17** | **-14** | **-11** | **-8** |
| AAC | 0.81 | 0.85 | 0.90 | 0.94 |
| ACP | 1.07 | 0.97 | 0.85 | 0.98 |
| CPG | 0.85 | 0.96 | 0.52 | 1.22 |
| **PGP** | 0.44 | 0.10 | **2.20** | 0.08 |
| GPY | 0.98 | 1.61 | 0.86 | 1.53 |
| PYA | 1.47 | 1.27 | 1.53 | 1.10 |
| YAC | 0.94 | 1.13 | 0.92 | 1.18 |
| PYR | 1.03 | 0.78 | 1.11 | 0.97 |

Cloned sequence: AAC**EPG**YAC**EPG**YAC**EPG**YR

| Ribosome pausing 5Pseq (log2 Fold change eIF5Ats/wt) | | | | |
|---|---|---|---|---|
| **Motif** | **Ribosome position** | | | |
| | **-17** | **-14** | **-11** | **-8** |
| AAC | 0.81 | 0.85 | 0.90 | 0.94 |
| ACE | 0.94 | 1.17 | 1.21 | 1.02 |
| CEP | 1.21 | 0.87 | 0.88 | 1.11 |
| **EPG** | 0.17 | -0.11 | 0.16 | **2.19** |
| PGY | 1.22 | 1.24 | 1.22 | 1.29 |
| GYA | 1.19 | 0.95 | 1.01 | 0.87 |
| YAC | 0.94 | 1.13 | 0.92 | 1.18 |
| GYR | 1.25 | 0.92 | 1.26 | 0.92 |

Cloned sequence: AAC**TQA**YAC**TQA**YAC**TQA**YR

| Ribosome pausing 5Pseq (log2 Fold change eIF5Ats/wt) | | | | |
|---|---|---|---|---|
| **Motif** | **Ribosome position** | | | |
| | **-17** | **-14** | **-11** | **-8** |
| AAC | 0.81 | 0.85 | 0.90 | 0.94 |
| ACT | 1.12 | 0.99 | 0.92 | 1.13 |
| CTQ | 1.06 | 0.91 | 1.13 | 0.88 |
| TQA | 0.06 | 0.06 | -0.17 | -0.28 |
| QAY | 0.96 | 1.40 | 1.08 | 1.11 |
| AYA | 0.88 | 0.80 | 1.15 | 0.79 |
| YAC | 0.94 | 1.13 | 0.92 | 1.18 |
| AYR | 1.19 | 0.61 | 0.86 | 0.67 |

**Figure EV1**

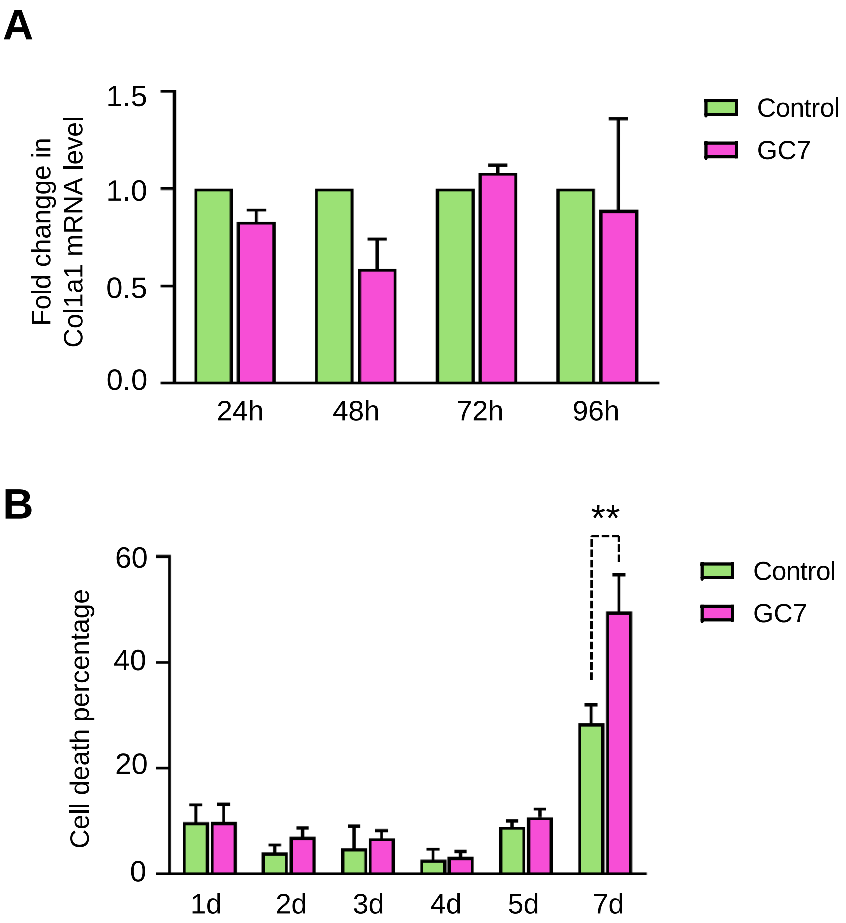
A
1.5
1.0
0.5
0.0
Fold changge in
Col1a1 mRNA level
24h
48h
72h
96h
Control
GC7
B
**
60
40
20
0
Cell death percentage
1d
2d
3d
4d
5d
7d
Control
GC7

**Figure EV2**

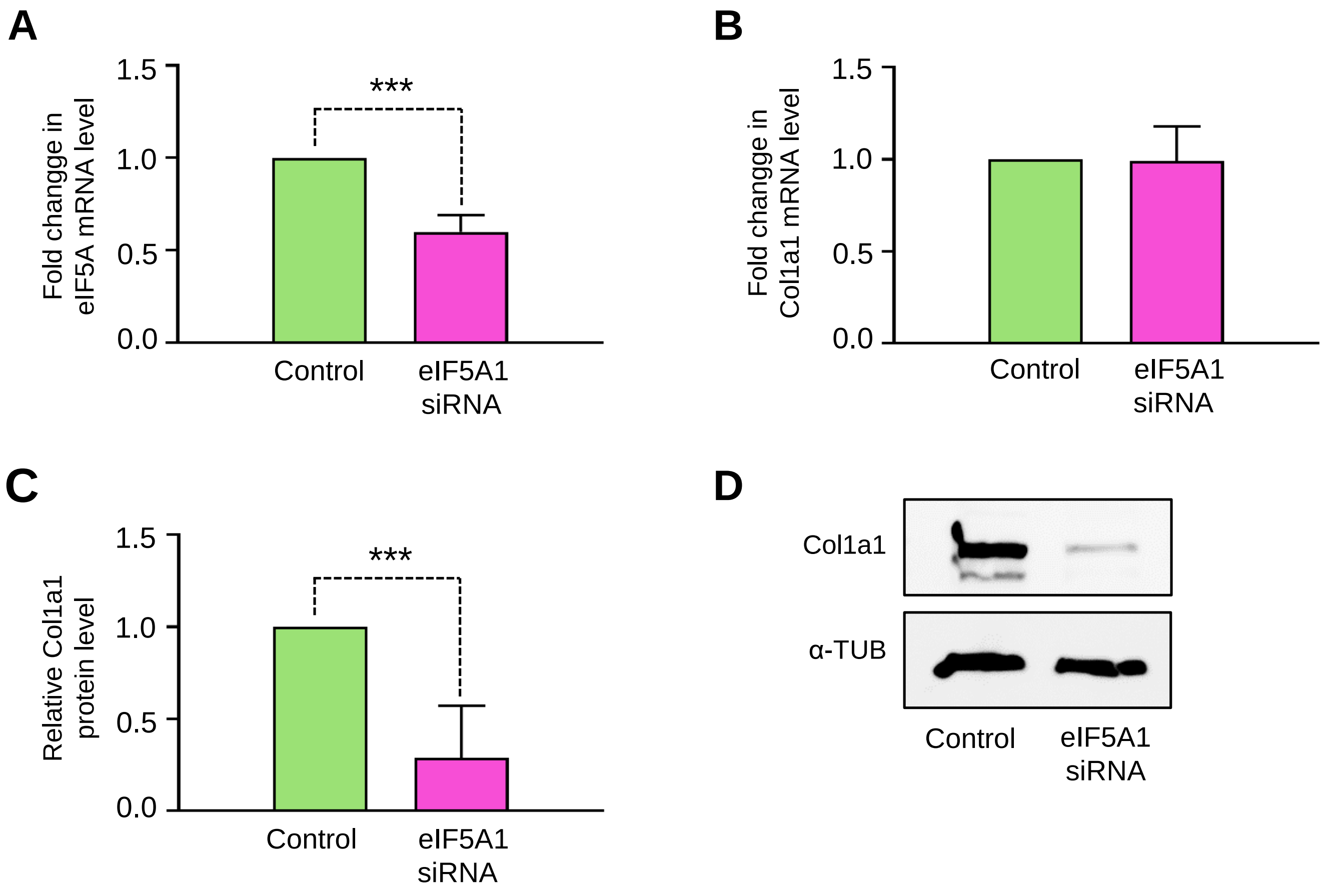
A
1.5
1.0
0.5
0.0
Fold changge in
eIF5A mRNA level
***
Control
eIF5A1
siRNA
B
1.5
1.0
0.5
0.0
Fold changge in
Col1a1 mRNA level
Control
eIF5A1
siRNA
C
1.5
1.0
0.5
0.0
Relative Col1a1
protein level
***
Control
eIF5A1
siRNA
D
Col1a1
α-TUB
Control
eIF5A1
siRNA

**Figure EV3**

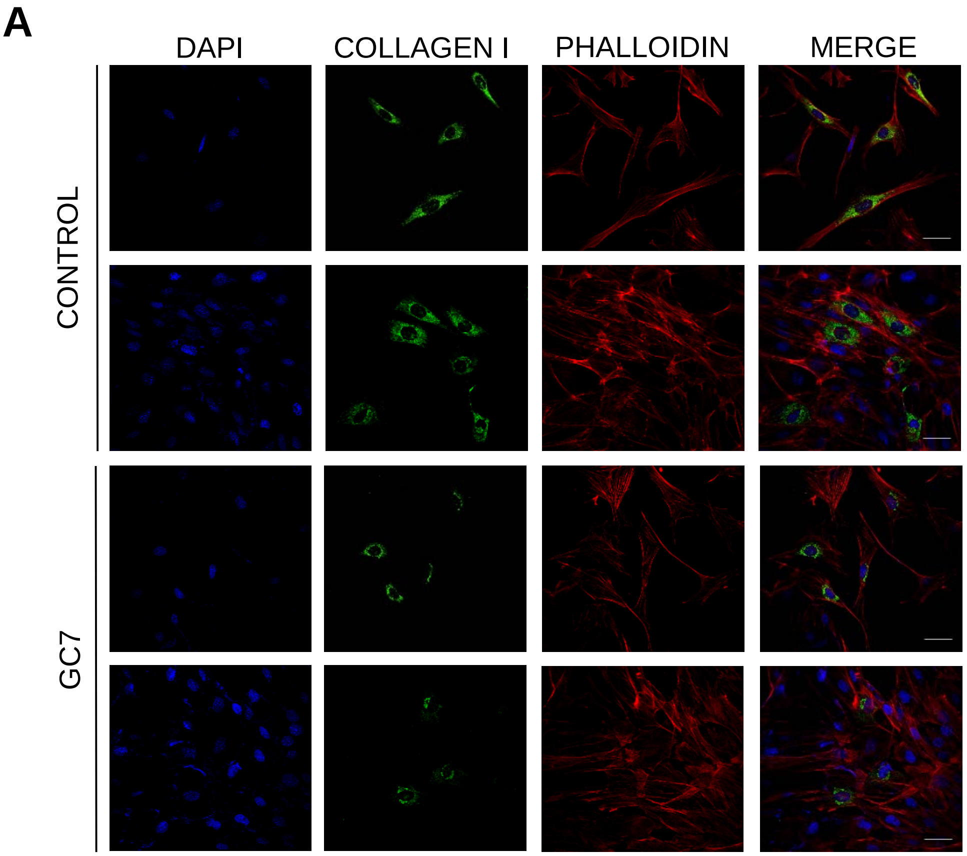


**Figure EV4**
