## Supplemental Tables S1 and S2 for "Hypusinated eIF5A is required for the translation of collagen"

**Appendix**

**Contents:**

Appendix Table S1

Appendix Table S2

**Appendix Table S1. Yeast strains used in this study**

| Name | Genotype | Source |
|---|---|---|
| **BY4741** | MATa *his3∆0 leu2∆0 met15∆0 ura3∆0* | Euroscarf |
| ***tif51A-1*** | MATa *his3∆0 leu2∆0 met15∆0 ura3∆0 tif51A-1::KanMX* | (Li *et al.*, 2011) |
| **J697** | MATα *trp1-Δ63 ura3-52 leu2-3 leu2-112 gcn2Δ tif51b::NAT tif51a::KANMX4 + p[TIF51A, LEU2]* | (Saini *et al.*, 2009) |
| **J699** | MATα *trp1-Δ63 ura3-52 leu2-3 leu2-112 gcn2Δ tif51b::NAT tif51a::KANMX4 + p[tif51a-S149P, LEU2]* | (Saini *et al.*, 2009) |

**Appendix Table S2. Oligonucleotides used in this study**

| Primer | Sequence (5'-3') |
|---|---|
| **mRNA levels assessed by RT-qPCR in mouse fibroblasts** | |
| ATF6-F | AGCTGTCTGTGTGATGATAG |
| ATF6-R | GTGATCATAGCTGTAACTGTC |
| BIP-F | CATGGTTCTCACTAAAATGAAGG |
| BIP-R | GCTGGTACAGTAACAACT |
| CHOP-F | TGTTGAAGATGAGCGGGTGG |
| CHOP-R | CGTGGACCAGGTTCTGCTTT |
| COLLAGEN I α1-F | GAAGCACGTCTGGTTTGG |
| COLLAGEN I α1-R | ACTCGAACGGGAATCCAT |
| eIF5A1-F | TGGTTTAGGTTCCCCTCTCC |
| eIF5A1-R | GCTTGGGGGTGAAGGACTAT |
| GAPDH-F | AGGTCGGTGTGAACGGATTTG |
| GAPDH-R | GGGGTCGTTGATGGCAACA |

| | |
|---|---|
| **Cloning into pCM179 yeast plasmid** | |
| COLLAGEN I-α1-F | ACGCAAACACAAATACACACACTAAATTACCGGATCAATTCGGGGATGTTCAGCTTTGTGGAC |
| COLLAGEN I-α1-R | CGACGTTGTAAAACGACGGCCAGTGAATCCGTAATCATGGTCATACCCTTGGGACCTGGAGG |
| BNI 1-F | ACGCAAACACAAATACACACACTAAATTACCGGATCAATTCGGGGATGTTGAATACAGATGGCGCAGAAGAT |
| BNI 1-R | CGACGTTGTAAAACGACGGCCAGTGAATCCGTAATCATGGTCATTTTCTTGTGTGGACGAGGATA |
| **Cloning into pDL202 yeast plasmid** | |
| **Primary sequences** | |
| Collagen IV-α1 | GCTGCTGCTGCTGCTTGTCCTCCGGGATTTACTGGACCACCGGGTCCTCCAGGCCCTCCTGGACCTCCTGGATACCGATGCGATCAAACG |
| 3PPP | GCTGCTGCTGCTGCTTGTCCACCACCATACGCTTGTCCACCACCATACGCTTGTCCACCACCATACCGATGCGATCAAACG |
| 3PGP | GCTGCTGCTGCTGCTTGTCCAGGTCCATACGCTTGTCCAGGTCCATACGCTTGTCCAGGTCCATACAGATGTGACCAAACT |
| 3EPG | GCTGCTGCTGCTGCTTGTGAACCAGGTTACGCTTGTGAACCAGGTTACGCTTGTGAACCAGGTTACAGATGTGACCAAACT |
| 3TQA | GCTGCTGCTGCTGCTTGTACTCAGGCTTACGCTTGTACTCAGGCTTACGCTTGTACTCAGGCTTACAGATGTGACCAAACT |
| **Secondary sequences** | |
| pDL202 5' | AAATCGTTCGTTGAGCGAGTTCTCAAAAATGAACAAATGTCGACGGCTGCTGCTGCTGCTTGT |
| pDL202/303 3' | TTCGAGCTCAGGGAAGTTGAAGGATCCGAACGTTTGATCGCATCGGTA |
